# Designer Fat Cells: Adipogenic Differentiation of CRISPR-Cas9 Genome-Engineered Induced Pluripotent Stem Cells

**DOI:** 10.1101/2023.10.26.564206

**Authors:** E. V. Ely, A. T. Kapinski, S. G. Paradi, R. Tang, F. Guilak, K. H. Collins

## Abstract

Adipose tissue is an active endocrine organ that can signal bidirectionally to many tissues and organ systems in the body. With obesity, adipose tissue is a source of low-level inflammation that contributes to various co-morbidities and damage to downstream effector tissues. The ability to synthesize genetically engineered adipose tissue could have critical applications in studying adipokine signaling and the use of adipose tissue for novel therapeutic strategies. This study aimed to develop a method for non-viral adipogenic differentiation of genome-edited murine induced pluripotent stem cells (iPSCs) and to test the ability of such cells to engraft in mice *in vivo*. Designer adipocytes were created from iPSCs, which can be readily genetically engineered using CRISPR-Cas9 to knock out or insert individual genes of interest. As a model system for adipocyte-based drug delivery, an existing iPSC cell line that transcribes interleukin 1 receptor antagonist under the endogenous macrophage chemoattractant protein-1 promoter was tested for adipogenic capabilities under these same differentiation conditions. To understand the role of various adipocyte subtypes and their impact on health and disease, an efficient method was devised for inducing browning and whitening of IPSC-derived adipocytes in culture. Finally, to study the downstream effects of designer adipocytes *in vivo*, we transplanted the designer adipocytes into fat-free lipodystrophic mice as a model system for studying adipose signaling in different models of disease or repair. This novel translational tissue engineering and regenerative medicine platform provides an innovative approach to studying the role of adipose interorgan communication in various conditions.

## Introduction

Obesity is a rapidly increasing public health condition (affecting about one-third of the world population (and is characterized by metabolic disturbance and pathological adipose signaling (Hildreth *et al*., 2021). Although adipose tissue was long considered to be inert, the active endocrine roles of adipocytes, a major cell type in adipose tissue, and other immune cell populations resident in adipose tissue have pleiotropic roles in both maintaining health as well as contributing to disease and reduced healthspan (Kahn *et al*., 2019; Kajimura, 2017). For example, low-level systemic inflammation from obesity results in metabolic dysfunction, inducing a milieu of obesity-linked diseases (Scarpellini and Tack, 2012) like diabetes mellitus (Singh *et al*., 2013), cardiovascular disease (Singh *et al*., 2013), and various musculoskeletal disorders (Anandacoomarasamy *et al*., 2008; Collins *et al*., 2018; Griffin and Guilak, 2008; King *et al*., 2013; Velasquez and Katz, 2010). These interactions have been difficult to disentangle due to interorgan crosstalk (Kirk *et al*., 2020; Velez *et al*., 2023) between adipose and downstream effector tissues (Kirk *et al*., 2020) but could be the key to understanding the onset and progression of diseases at the interface of aging and obesity or in multi-organ aging.

Adipose tissue exists in three phenotypically and functionally distinct subtypes – white adipose tissue (WAT), brown adipose tissue (BAT), and beige adipose tissue (BeAT) — which are distributed heterogeneously throughout the body in anatomical regions called depots (Marcadenti and de Abreu-Silva, 2015). In obesity, there is an overabundance of WAT; however, the extent of metabolic dysfunction varies depending on the location (e.g., visceral vs. subcutaneous) and adipokine expression levels of adipose depots (Fried *et al*., 1998; Marcadenti and de Abreu-Silva, 2015). Adipokines mediate inflammation and regulate systemic metabolic, skeletal, and reproductive processes (Griffin *et al*., 2009; Kwon and Pessin, 2013), but when dysregulated, adipokines can directly influence many obesity-associated diseases (Fasshauer and Blüher, 2015; Unamuno *et al*., 2018). Some examples of adipokines include adiponectin, leptin, visfatin, resistin, interleukin 6 (IL-6), and tumor necrosis factor alpha (TNF-α) (Kwon and Pessin, 2013). Conversely, BAT is considered a potential therapeutic and metabolically healthy tissue associated with improved caloric expenditure and thermal regulation (Marcadenti and de Abreu-Silva, 2015; Nicholls and Locke, 1984). An improved understanding of adipose tissue’s physiologic and pathologic roles will provide new insights toward controlling and leveraging adipose signaling for novel disease therapeutics in various disease states (Batún-Garrido *et al*., 2018; Grewal and Buechler, 2023; Recinella *et al*., 2020; Su *et al*., 2019) (i.e., COVID-19, cardiovascular disease, diabetes, cancer, metabolic syndrome). It may be similarly possible to harness adipose tissue phenotypes – such as BAT in the studies mentioned above – for enhanced therapeutic effects. However, the mechanisms by which specific adipose tissue subtypes and their unique signaling attributes affect disease progression are unknown, primarily due to a lack of effective adipose tissue models.

Numerous animal models have been utilized to study the effects of obesity, largely involving single gene knockout or transgenic mice and/or obesogenic diets to study metabolic changes (Coleman, 1978; Collins *et al*., 2015; Griffin *et al*., 2010; Griffin and Guilak, 2008; Griffin *et al*., 2009; Hariri and Thibault, 2010; Levin and Dunn-Meynell, 2002). More recently, fat-free lipodystrophic (LD) mice, coupled with implanted fat pads or mouse embryonic fibroblasts (MEFs) have been used to study the effects of different aspects of adipose signaling on musculoskeletal health (Collins *et al*., 2022; Collins *et al*., 2021). However, the heterogeneity of MEFs can result in the restoration of circulating adipokines that are known to contribute to pro-inflammatory catabolic activities and metabolic complications (Collins *et al*., 2021; Ferguson *et al*., 2018) (i.e., IL-6). Due to the inability to tightly control MEF signaling, evaluating induced pluripotent stem cells (iPSCs) as a cell source was a logical step to precisely engineer adipocytes.

iPSCs have multiple advantages as a source for developing “designer” cells, as single clonal iPSCs can be genome-edited, expanded, and differentiated widely into a variety of cell and tissue types (Diekman *et al*., 2012; Zomer *et al*., 2015). iPSCs have been successfully differentiated into adipocytes by overexpressing *Ppar*γ using adenovirus vector-mediated transient transduction (Mohsen-Kanson *et al*., 2014; Tashiro *et al*., 2009) and pharmacological overexpression using rosiglitazone or troglitazone (Fayyad *et al*., 2019; Kim *et al*., 2014; Tchoukalova *et al*., 2000). Developing a streamlined, virus-free protocol for adipogenesis allows users to generate large numbers of designer adipocytes from single clonal iPSCs consistently without concerns of losing differentiation potential. This reproducible method can be easily scaled up for clinical translation. This proposed model of designer adipocytes can be implanted into animal models of adipose ablation (Collins *et al*., 2021; Huang-Doran *et al*., 2010; Klaus *et al*., 1998; Savage, 2009), like lipodystrophic mice, to directly disentangle the role of adipokine signaling in downstream signaling, homeostasis, and disease progression.

The overall goal of this study was to use iPSCs to genome-engineer designer adipose tissue to serve as a regenerative medicine platform to unravel the roles of adipokines in disease progression. Initially, a protocol was created to differentiate adipocytes from murine iPSCs in a virus-free manner. Four phenotypes of designer adipocytes were generated, spanning mechanistic and drug delivery applications. First, CRISPR-Cas9 genome engineering was used to create a leptin knockout iPSC line, as leptin is an adipokine and pro-inflammatory mediator that is consistently increased with obesity and downstream catabolic tissue effects (Krempler *et al*., 1998; Perakakis *et al*., 2021). Designer adipocytes provide a tool to allow us to disentangle the role of leptin (Collins *et al*., 2022) or other adipokines in regulating effector tissues. Next, designer adipocytes were developed to deliver anti-cytokine therapies, to serve as a novel adipose tissue-based drug delivery strategy. For proof of concept, an existing therapeutic iPSC line was used that has been rewired to inhibit interleukin-1 (IL-1) signaling by endogenously producing interleukin-1 receptor antagonist (IL-1Ra) in the presence of inflammation (Brunger *et al*., 2017a; Brunger *et al*., 2017b).

Lastly, we propose to use designer adipose as a tool to recapitulate and decode the mechanistic and phenotypic differences between WAT and BAT *in vitro* using two protocols to address the unmet need for engineered brown and white adipose tissues. Previous studies have utilized CRISPR-Cas9 technology to generate human brown-like adipocytes from ASCs derived from subcutaneous white adipose tissue, which improved glucose tolerance, insulin sensitivity, and energy expenditure upon transplantation (Tsagkaraki *et al*., 2021; Wang *et al*., 2020). This work focused on creating an efficient approach for the browning and whitening of adipocytes from iPSCs. As browning of fat has been suggested as a therapeutic opportunity, we are interested in exploring designer brown adipocytes in the context of obesity-linked disorders and regenerative applications.

## Results

### PDiPSCs Demonstrate Adipocyte Phenotype After Virus Free Culture

A virus-free protocol was developed for PDiPSCs by based on a commercially available adipogenic media designed for MSC adipogenesis. We then compared this protocol to published methods using lentiviral transduction of *Ppar*γ, a transcription factor integral for the differentiation of mesenchymal-like cells to adipocytes. Increasing concentrations of the virus were tested, and 50μL of the virus, or the high virus dose, displays the highest concentration of rounded, lipid-like cells, while 30μL of the virus, or the low dose, yields the second-highest concentration of lipid-like cells [Figure 1a]. Oil Red O (ORO) staining of samples at day 14 with different (no, low, and high) virus concentrations and media demonstrated differentiation of lipid-containing cells using expansion media for both the low and high doses of virus, adipogenic media with both low and high doses of virus, and adipogenic media with no virus [Figure 1b]. For the virus conditions, the highest virus concentration [50μL of the virus] yielded the highest density of PDiPSCs stained with ORO in both expansion and adipogenic media conditions. In contrast, the low virus condition resulted in fewer lipid-containing cells overall. As such, the highest concentration virus dose was used as a control to confirm PDiPSC-adipocyte differentiation in a virus-free manner. Under 20X magnification, the adipogenic media group with no virus, expansion media group with virus, and the adipogenic media group with no virus revealed cells with a rounded, plump morphology in clusters of 3-4 cells, and ringlets of lipids were plentiful. The expansion group with virus also demonstrated a rounded morphology, but fewer clusters were observed compared to the adipogenic media group [Figure 1c].

**Fig. 1.**
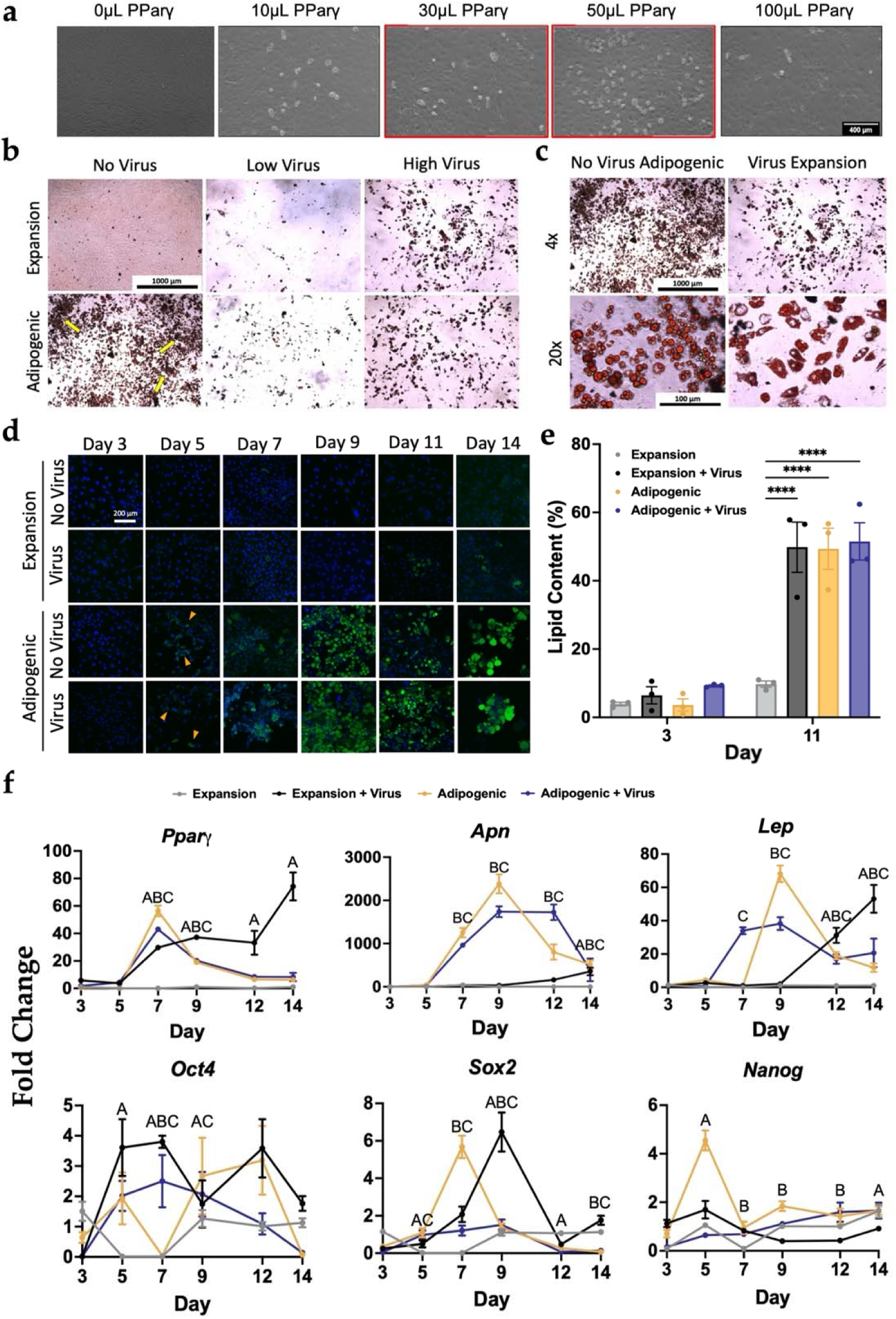
PDiPSC adipogenesis using either viral antivirus-free methods, (a) WT PDiPSCs were cultured with varying concentrations of *Pparγ* virus. After 14-14-days culture in expansion media with virus, 50 pL of *Pparγ* virus displays the highest concentration of adipocytes like cells. WOpL of virus resulted in less cell density Scale bar is 400pm. All groups received 5μL puromycin WT PDiPSCs were then cultured with no *Pparγ* virus low dose of *Ppar/’* virus (3.125pl/cm^2^), or high dose *Pparγ* virus (5.208pl/cm^2^) in expansion or adipogei media for 14 days, (b) All groups were stained with (Red O and imaged at 4x. All groups were stained with Oil Red O stain, where the red stain indicates lipids content in cells. Scale bar is 1000pm (c) 20x images virus-free adipogenic media and virus-expansion media groups were taken to examine morphology of the PDiPSCs. Scale bars are 1000pm and 100pm. (d) WT PDiPSCs with no virus and virus were cultured expansion and adipogenic media conditions for 14 da and stained with BODIPY/DAPI. BODIPY, green, stained for lipids and DAPI, blue, stains nuclei. Scale bar 200pm. (e) Lipid content of the PDiPSCs was measured at days 3 and 11 using the BODIPY images. Ba represent lipid content (%) ± SEM *(n=3). Asterisks* represent significance (**** *p<.00005)* compared with the WT PDiPSCs expansion media control, (f) Cells from groups above were collected at various timepoints I gene expression characterization. *Ppary, Apn,* and *L Oct4,Sox2,* and *Nanog* were target genes evaluated, we target genes evaluated. Values represent fold change SEM (n=3). Letters represent significance *(A p<0 expansion media + virus; B p<.05 adipogenic media;* C *p<.05 adipogenic media + virus).* All statistics were run using a way ANOVA with Sidak’s post-hoc test. SEM, standard error of the mean.

A time course experiment was used to evaluate PDiPSC adipogenesis with and without the virus. Cells were fixed 3-, 5-, 7-, 9-, 11-, and 14-days after plating and stained with BODIPY/DAPI [Figure 1d]. In the expansion media groups, few cells stained positive from days 3-11. In the expansion media condition with the virus, some BODIPY stain was evident on day 14. In the adipogenic media groups, BODIPY staining was present as early as day 5 in the virus and no-virus groups. The density of BODIPY-stained PDiPSCs illustrated a trend toward increased staining in both adipogenic media groups with time up to day 14.

Image analysis tools were used to quantify prior observations with the BODIPY time course images to quantify lipid content per cell in each image [Figure 1e]. Lipid content was quantified as the percentage of lipid content per cell. For this, PDiPSCs cultured in expansion or adipogenic media with virus or no virus were compared on day 3 and day 11. Results of these analyses revealed no significant differences between all treatment groups from the control expansion media condition at day 3 (expansion with virus 6.4 ± 2.5%, adipogenic no virus 3.6 ± 1.8%, adipogenic with virus 9.3 ± 0.2%). However, by day 11, all groups demonstrated significantly increased (*p<0.001*) percent lipid content than the control (expansion with virus 49.9 ± 7.4%, adipogenic no virus 49.4 ± 6.0%, adipogenic with virus 51.5 ± 5.5%). Together, these data indicated no overt benefit to using virus compared to adipogenic media to differentiate PDiPSCs into adipocytes based on morphological presentation.

### PDiPSCs Demonstrate Increased Expression of Adipogenic Genes in Virus Free Culture

RT-qPCR studies were performed to explore differences in mRNA expression in virus and no virus expansion media and adipogenic media conditions over a 14-day time course. Transcriptional changes during adipogenesis were measured by evaluating pluripotency and adipogenic marker gene expression in cultured PDiPSCs.

*Ppar*γ mRNA levels were significantly increased in all groups on day 7 and 9 compared to control expansion media (*p<0.001*) [Figure 1f]. At day 11 and 14, only the expansion media with virus group had significantly increased expression when compared to mRNA levels measured in the control group (*p<0.001*). Strikingly, peak *Ppar*γ expression for the adipogenic media condition was at day 7 (56.4 ± 3.9 fold change), suggesting that PDiPSCs differentiated into adipocytes by the later time points. We next evaluated adiponectin, *Apn*, which is highly expressed by mature adipocytes (Farmer, 2005), indicating commitment to adipogenesis. Adipogenic media groups with virus and no virus exhibited significantly higher mRNA levels of *Apn* starting at day 7 (*p<0.001*), with peak fold changes observed at day 9 for both (no virus 2384.0 ± 221.6 fold change, with virus 1737 ± 125.9 fold change). These outcomes were consistent with the peak expression of *Ppar*γ at day 7, which would precede commitment to adipocytes. Moreover, the expansion media with virus group had significantly increased *Apn* expression compared to expansion media control only at day 14 (*p<0.001*), suggesting that adipocyte differentiation occurred over varying time courses with different methods. Lastly, mRNA levels for leptin, *Lep*, whose circulating expression is proportional to the amount of fat in the body or the number of adipocytes present (Krempler *et al*., 1998; Perakakis *et al*., 2021), were evaluated. The adipogenic media group with virus exhibited significantly higher mRNA levels of *Lep* compared to the expansion media control by day 7 (*p<0.001*), while the adipogenic group with no virus was significantly different from control and had peak mRNA expression at day 9 (68.2 ± 4.96 fold change*, p<0.001*). The expansion media with virus condition peak *Lep* mRNA levels were observed at day 14 (*p<0.001*). These findings were consistent with the trends observed in the other markers for adipogenesis. Taken together, mRNA fold changes for all three genes, *Ppar*γ*, Apn*, and *Lep,* [Figure 1f] were significantly increased using a virus-free approach compared to expansion media control groups as early as 5 days in culture. mRNA levels of key pluripotency markers octamer binding transcription factor 4 (*Oct4*), SRY-box transcription factor *(Sox2*), and nanog homeobox (*Nanog*) confirmed that the PDiPSCs committed to an adipogenic cell lineage and did not de-differentiate back into iPSCs from their mesenchymal-like PDiPSC state [Figure 1f].

### CRISPR-Cas9-Edited Leptin Knockout Adipocytes

To establish a platform for modifying adipokine genes with CRISPR-Cas9 and systematically testing the impact of secreted factors from engineered designer adipocytes, we initially employed engineered leptin knockout (KO) adipocytes as a proof of concept. We hypothesized that PDiPSCs with a leptin knockout would display robust adipogenesis following the validated virus-free culture protocol. Leptin KO cells grown in expansion and adipogenic media showed no positive staining with ORO, indicating no measurable presence of lipids. Days 5, 9, and 14 are shown as references [Figure 2a]. To confirm this finding, BODIPY fluorescent staining for lipids was conducted, which similarly did not reveal any minor positive staining until day 11 and day 14 in the adipogenic media group [Figure 2b]. Consistent with previous reports of leptin’s role in cell differentiation (Scheller *et al*., 2010), leptin KO cells proliferated rapidly and at times, were overconfluent, and lifted off the bottom of the culture well using the typical cell seeding conditions. To better understand the lack of adipocyte-like morphology, mRNA levels of key adipogenic genes were measured [Figure 2c]. WT PDiPSCs in adipogenic media had significantly higher (*p<0.001*) gene expression compared to the expansion media control, while surprisingly, the two leptin KO conditions had significantly lower (*p<0.001*) *Ppar*γ mRNA levels compared to control. WT PDiPSCs in adipogenic media had significantly higher mRNA levels (*p<0.001*) of *Apn* compared to the control, while the leptin KO PDiPSCs were similar to control. mRNA levels of *Lep* were not detected in leptin KO PDiPSCs. Together, these data suggest leptin is required for PDiPSC adipogenic differentiation, and as such, this hypothesis was directly tested next.

**Fig. 2.**
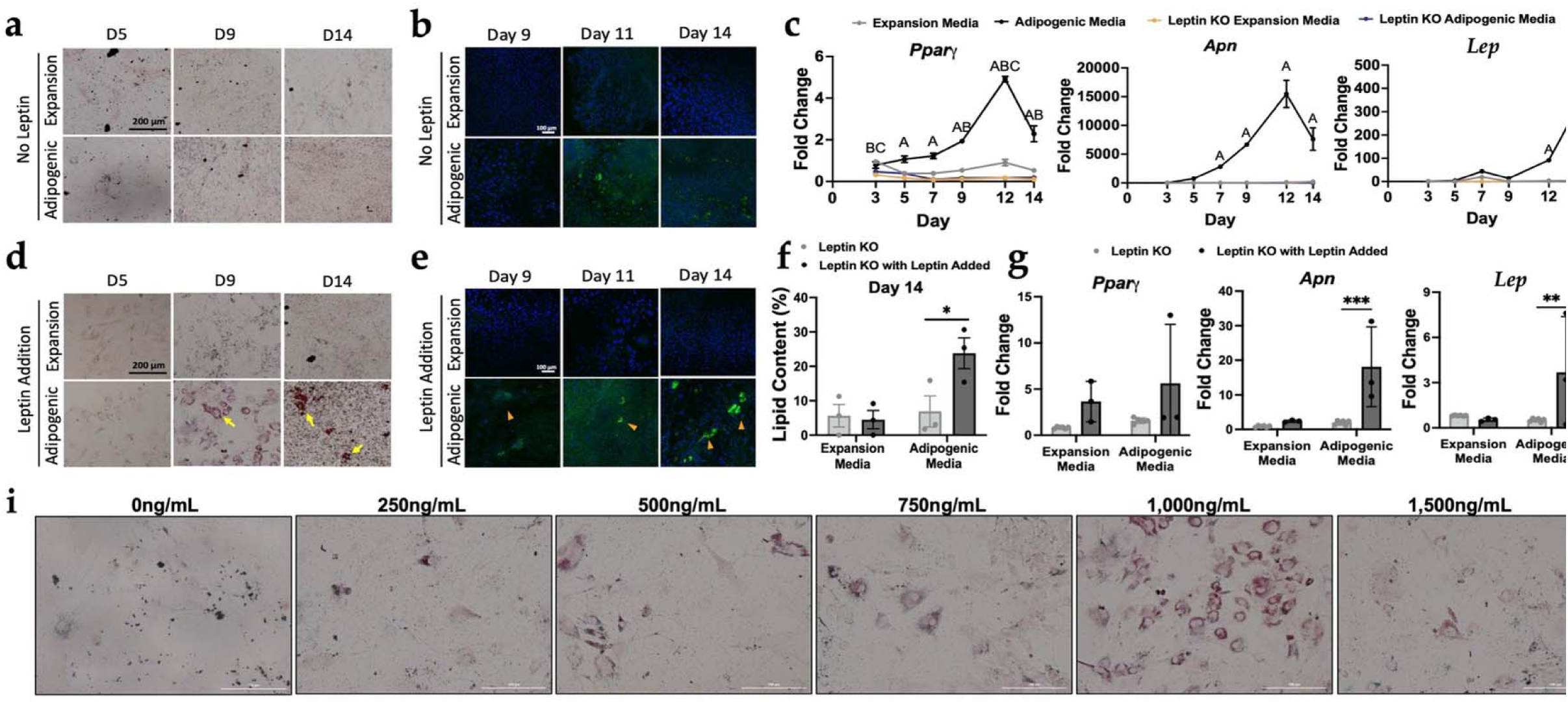
Leptin knockout PDiPSCs are capable of adipocyte differentiation, but not without supplemental leptin. Leptin KO PDiPSCs were cultured in expansion adipogenic media for 14 days, (a) All groups were stained with Oil Red O and Days 5, 9, and 14 are shown here. Red stain indicates lipid content in cells. The yell arrows point to lipid-containing cells. Scale bar is 200pm. (b) All Groups were stained with BODIPY/DAPI BOD1PY, green, stains for lipids and DAPI, blue, stains nuc Scale bar is 100pm. (c) WT and leptin KO PDiPSCs were collected at various timepoints for gene expression characterization. *Ppary, Adip,* and *Lep,* were target gei evaluated. Values represent fold change + SEM (n=3). Letters represent significance (A *p<.05, WT adipogenic media;* B *p<.05 leptin KO expansion media; C p<.05 leptin, adipogenic media’)* from WT PDiPSCs expansion media control. In an effort to restore adipogenic differentiation, murine leptin was added to the culture media a supplement, (d) Leptin KO PDiPSCs with leptin added were cultured in expansion or adipogenic media and for 14 days and were stained with Oil Red O and days 5 and 14 as shown here. Scale bar is 200pm. (e) The same groups as (d) were stained with BODIPY/DAPI. Scale bar is 100pm. (f) Lipid content of the PDiPSCs (leptin 1 and leptin KO with leptin added) was measured at day 14 using the BODIPY images. Bars represent lipid content (%) ± SEM *(n=3). Asterisks* represent significance *p<0.01)* compared with the leptin KO PDiPSCs in adipogenic media, (g) Cells from leptin KO with and without leptin added were collected at various timepoints for g< expression characterization, day 9 is shown here. *Ppary, Apn,* and *Lep,* were target genes evaluated. Values represent fold change ± SEM (n=3). *Asteriks* repress significance (** *p<0.0001, *** p<0.0000T)* compared with the leptin KO PDiPSCs expansion media control, (i) For 9 days, Leptin KO PDiPSCs were treated with adipoge media and supplemented with varying doses of leptin. Samples were fixed and stained with Oil Red O. l,000ng/mL of leptin added to culture media displays the high concentration of lipid-containing cells. Scale bar is 200pm. All statistics were run using a 2-way ANOVA with Sidak’s post-hoc test. SEM, standard error of the mean.

### Leptin Supplementation in Culture Media Rescues Adipogenesis in Leptin Knockout iPSCs

To restore the typical growth kinetics of PDiPSCs with adipogenesis media, leptin was added to the cell culture media. Using the same measures of morphology and gene expression as above, restoration of adipogenesis was observed in leptin KO PDiPSCs with leptin added. At days 9 and 14, leptin KO PDiPSCs with leptin added in adipogenic media revealed positive lipid staining by ORO [Figure 2d]. Similarly, BODIPY staining revealed adipocytes in culture, demonstrated by green stain and surrounding blue DAPI stained nuclei [Figure 2e). Using the BODIPY images, percent of lipid content per cell was quantified for the leptin KO groups with and without leptin added [Figure 2f]. Leptin KO PDiPSCs with leptin added in adipogenic media had a significantly higher lipid content compared to leptin KO PDiPSCs with no leptin added (no leptin 6.9 ± 4.5%, leptin added 23.8 ± 4.5%, *p=*0.028). qPCR profiles were measured at day 9 and the leptin KO with leptin added group was compared to leptin KO samples for *Ppar*γ*, Apn*, and *Lep* gene expression [Figure 2g]. *Ppar*γ mRNA levels were not significantly increased when leptin was added to the culture media (expansion media *p=0.286,* adipogenic media *p=0.112*). However, *Apn* and *Lep* mRNA levels were significantly increased in the leptin KO adipogenic condition with leptin added (*Apn p<0.001*, *Lep p=0.007*). Taken together with the morphological outcomes in this cell line, adding leptin back into the system rescued adipogenesis. Varying leptin concentrations were tested to determine the optimal dose to recover adipogenesis in the leptin KO PDiPSCs, and 1,000ng/mL results in the highest density of lipid-laden cells, as demonstrated by ORO [Figure 2i].

### Designer Adipocytes as an Anti-Cytokine Therapy to Mitigate Inflammation

Using self-regulating iPSCs developed in a previous study (Brunger *et al*., 2017a; Brunger *et al*., 2017b; Choi *et al*., 2021; Pferdehirt, 2019; Pferdehirt *et al*., 2022), we developed engineered adipocytes capable of producing interleukin-1 receptor antagonist (IL-1Ra) in the presence of low-level inflammation. We tested the ability of Ccl2-IL1Ra PDiPSCs (IL-1Ra) to differentiate into adipocytes using the previously validated protocol and to maintain the ability to produce IL-1Ra post-differentiation. IL-1Ra PDiPSCs had similar morphology compared to WT PDiPSCs in culture, as shown in the ORO and BODIPY figures [Figures 3a-b]. In the ORO images, IL-1Ra and WT PDiPSCs demonstrated rounded lipid droplets at day 5. The number of lipid-containing cells increased with time in culture, indicated by the red staining [Figure 3a]. Between days 9-14, peak adipogenesis was observed by morphological and mRNA readouts for both groups. The BODIPY with DAPI counterstain illustrated that the IL-1Ra PDiPSCs in adipogenic media differentiated slightly earlier than the WT PDiPSCs at day 5, but both groups in adipogenic media displayed consistent adipogenesis through day 14 [Figure 3b]. Lipid content was quantified at days 3,9, and 14, and WT and IL1Ra groups in expansion and adipogenic media were compared to the control WT expansion media group [Figure 3c]. At day 3, all groups had similar mean percentage of lipid content (WT adipogenic 1.80 ± 0.84%, IL-1Ra expansion 2.9 ± 1.8%, IL-1Ra adipogenic 3.5 ± 1.9%). By day 9, there were significant differences in the WT and IL-1Ra adipogenic media groups. The WT PDiPSCs in adipogenic media mean lipid content was 18.2 ± 1.4% while the IL-1Ra PDiPCSs in adipogenic media had a mean lipid content of 17.9 ± 5.3% with a wide range of values from 12.5-28.5%. These data suggest that the IL-1Ra PDiPSCs may differentiate into adipocytes less efficiently under these conditions but can differentiate along the 14-day time course.

**Fig. 3.**
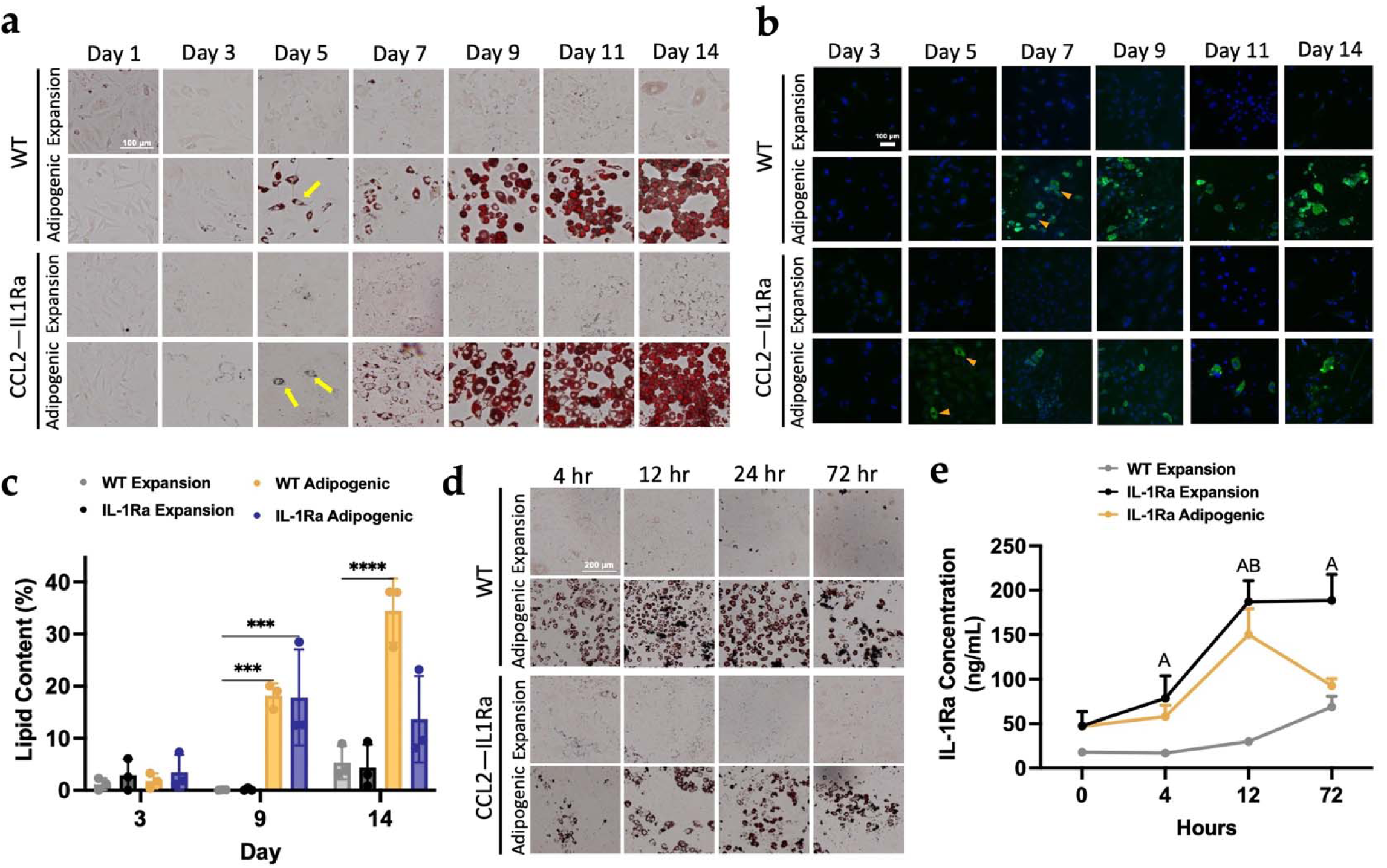
Adipogenic differentiation of genome-edited self-regulating anti-inflammatory iPSCs. WT PDiPSCs and Ccl2-ILlRa PDiPSCs were cultured in expansion or adipogenic media for 14 days and samples were fixed every other day from day 1 to day 14. (a) All groups were stained with Oil Red O stain, where the red stain indicates lipid content in cells. The yellow arrows point to lipid-containing cells. Scale bar is 200pm (b) WT and Ccl2-ILlRa PDiPSCs were stained with BODIPY/DAPI at various timepoints. BODIPY, green, stains for lipids and DAP1, blue, stains nuclei. Scale bar is 100 pm (c) Lipid content of the PDiPSCs was measured at days 3, 9, and 14 using the BODIPY images. Bars represent lipid content (%) ± SEM *(n=3). Asteriks* represent significance (*** *p<.0005, **** p<.00005)* compared with the WT PDiPSCs expansion media control. At Day 14, WT and Ccl2-ILlRa PDiPSCs were dosed with an inflammatory challenge to test the self-regulatory circuit, (d) WT and PDiPSCs were fixed 4-, 12-, 24-, and 72-hours after inflammatory challenge and stained with Oil Red O. (e) An ELISA was performed to determine the production of ILl-Ra protein using media from various timepoints of the inflammatory challenge. Values represent mean + SEM *(n=3).* Letters represent significance (A *p<.05, ILlRa+expansion media; B p<.05 ILlRa+adipogenic media).* All statistics were run using a 2-way ANOVA with Sidak’s post-hoc test. ELISA, enzyme-linked immunosorbent assay; SEM, standard error of the mean.

Ccl2-IL1Ra PDiPSCs were then analyzed to determine if the designer adipocytes produced IL-1Ra to similar levels of cells in expansion media in the presence of low-level inflammation. WT and IL-1Ra PDiPSCs were challenged with IL-1α and assessed for morphological and protein production changes. At the various timepoints, WT and IL-1Ra PDiPSCs stained positively for ORO, indicating that the cells persisted in culture throughout the inflammatory treatment [Figure 3e]. Protein production of IL-1Ra was quantified for each group along the 72-hour time course and compared to the control WT PDiPSCs in expansion media at each timepoint. Twelve hours after the inflammatory challenge, the IL-1Ra PDiPSCs produced a significantly higher amount of IL-1Ra (expansion media 188.0 ± 23.6 ng/mL, *p<0.001,* adipogenic media 150.0 ± 28.9 ng/mL, *p<0.001*) compared to the control (29.9 ± 2.5 ng/mL) [Figure 3f]. At 72 hours, the IL-1Ra PDiPSCs in expansion media again produced a significantly higher amount of IL-1Ra (188.0 ± 29.1 ng/mL, *p<0.001*) compared to the control, but the IL-1Ra PDiPSCs in adipogenic media did not produce a significantly higher amount compared to the control (92.6 ± 7.9 ng/mL, *p=0.577.* Therefore, rewired Ccl2-IL1Ra PDiPSCs designer adipocytes can produce anti-inflammatory mediators in the presence of an inflammatory environment.

### iPSC-Derived Brown and White Adipocytes

In this study, we aimed to develop brown and white adipose cell models in which adipokine expression can be precisely and efficiently edited using CRISPR-Cas9 to address key mechanistic gaps regarding the role of fat on disease pathophysiology. We hypothesized that PDiPSCs fed AdipoQual BAT or WAT media for white and brown adipogenesis would be capable of differentiating into distinct populations of brown and white adipocytes in culture. Morphology of the brown and white PDiPSCs and their gene expression profiles were analyzed. ORO images taken from day 0 to day 7 illustrated that the brown and white PDiPSCs behaved similarly to WT PDiPSCs, where lipid droplets appeared in culture by day 3 [Figure 4a]. By day 7, the WT and Brown PDiPSCs qualitatively appeared to have the highest number of lipid-containing cells but were closely followed by the white PDiPSCs. These initial data demonstrated that the AdipoQual BAT and WAT media were capable of inducing adipogenesis. Next, samples were stained with BODIPY and DAPI to determine if the lipid containing cells observed in ORO staining were adipocytes [Figure 4b]. Initial signs of adipocytes were spotted on day 3, but unlike the Oil Red O, the BODIPY images indicated fewer lipid-containing cells. The brown PDiPSCs differentiated into more adipocytes than the white PDiPSCs, which was consistent with the ORO findings. Differentiation efficiency and percent lipid content were quantified using the BODIPY confocal images [Figure 4c]. WT PDiPSCs in adipogenic media, brown PDiPSCs, and white PDiPSCs were compared to control WT PDiPSCs in expansion media at day 0 and day 7. Of note, at day 7, the WT adipogenic (0.44 ± 0.05, *p<0.001*) and brown PDiPSCs (0.18 ± 0.04, *p<0.007*) had significantly higher differentiation efficiencies compared to the WT control. Although not statistically significant, there was a trend toward increased differentiation efficiency of the white PDiPCs (0.11 ± 0.05, *p<0.*145) compared to the control.

**Fig. 4.**
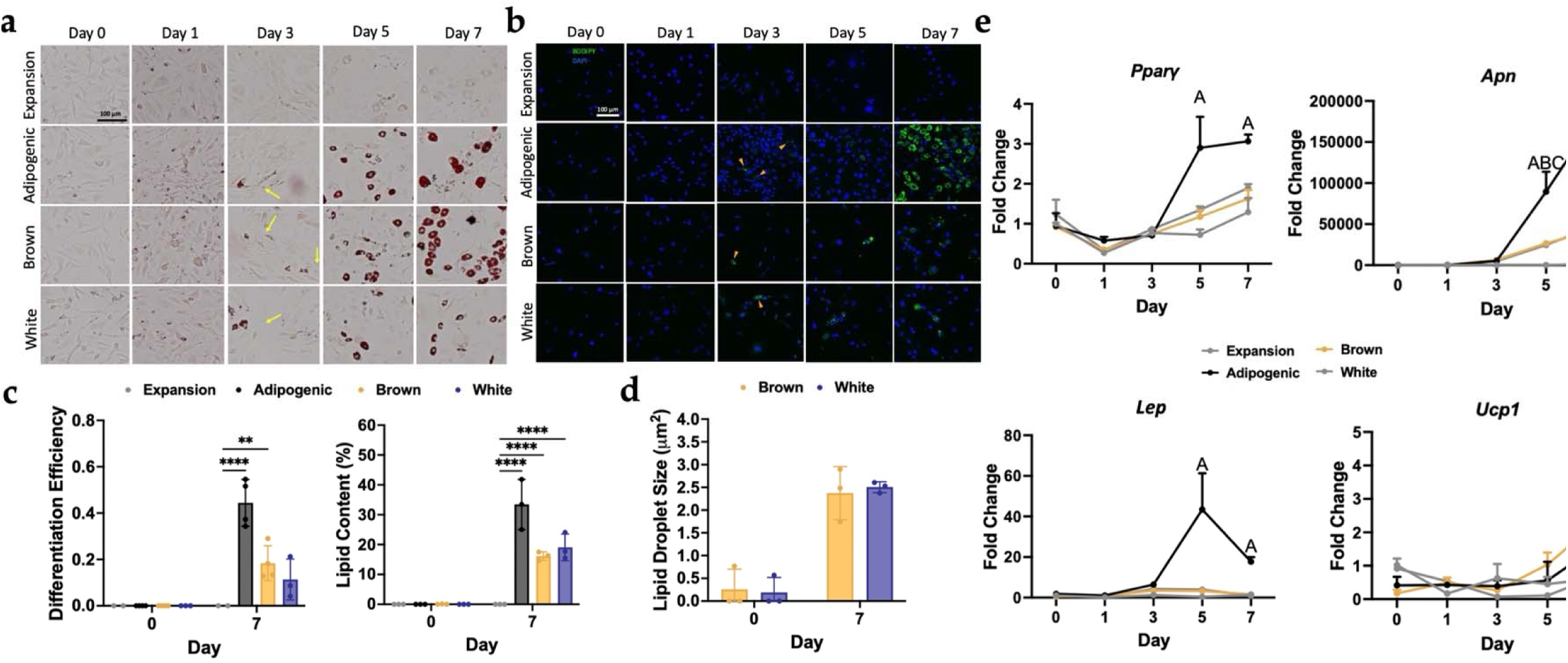
Brown and White PDiPSCs display adipocyte-like morphology. WT PDiPSCs were cultured with various media (expansion, adipogenic, brown, and white days, (a) All groups were stained with Oil Red O stain, where the red stain indicates lipid content in cells. The yellow arrows point to lipid-containing cells. Scale 100pm. (b) All groups were stained with BODIPY/DAPI at various timepoints. BODIPY, green, stains for lipids and DAPI, blue, stains nuclei. Scale bar is lOOp Differentiation efficiency and lipid content of all groups was measured at days 0 and 7 using the BODIPY images. Bars represent differentiation efficiency or lipid c< (%) ± SEM (n=3). *Asteriks* represent significance C’ *jx.005, *** p<.0005,* **** p<.00005) compared with the WT PDiPSCs expansion media control, (d) Lipid droplet siz measured at Days 0 and 7. Bars represent lipid droplet size + SEM (n=3). *Asteriks* represent significance (** *p<.005, *** p<.0005,* **** p<. 00005) compared with tb PDiPSCs expansion media control, (e) Cells from all groups above were collected at various timepoints for gene expression characterization. Ppar/, *Apn, Lep,* and were target genes evaluated. Values represent fold change ± SEM *(n=3).* Letters represent significance (A *p<.05, adipogenic media; B p<.05 brown media;* C *p<.05 white i* All statistics were run using a 2-way ANOVA with Sidak’s post-hoc test. SEM, standard error of the mean.

Key differentiating characteristics between different adipocyte subtypes are lipid size and number. Lipid size of the brown and white PDiPSCs were quantified with the hypothesis that the brown PDiPSCs will have smaller lipid sizes than the white PDiPSCs. Lipid size at day 0 and day 7 was evaluated, and there were no significant changes in lipid size amongst any of the groups [Figure 4d]. Brown adipocytes at day 7 had a mean lipid droplet size of 2.4±0.33μm and white adipocytes had a mean lipid size of 2.50 ±0.07μm. We hypothesize that at later time points, we will see a more robust difference between the two groups, especially given the wide range of lipid sizes for brown PDiPSCs and tight sample range for white PDiPSCs.

Lastly, selected adipogenic markers were analyzed to investigate whether the brown and white PDiPSCs displayed adipogenic phenotypes and, most importantly, whether the brown PDiPSCs were characterized by *Ucp-1* expression, the defining marker for brown adipose [Figure 4e]. Only the WT adipogenic group had significant changes in gene expression for *Ppar*γ *(p<0.001*) when compared to the WT expansion media control. The brown and white PDiPSC groups were not significantly different from the control group from day 0 to day 7. All groups had significantly increased *Apn* expression compared to the control (*p<0.001*) by days 5 and 7. *Lep* mRNA levels were significantly increased in the WT adipogenic group compared to control (*p<0.021*) on day 7. Excitingly, mRNA levels of *Ucp-1* were increased at day 7 in the brown PDiPSC group compared to the control (*p<0.048*). These data suggest that, in addition to white adipocytes, brown UCP-1 producing adipocytes can be differentiated from iPSCs.

### iPSC-Derived Adipocytes have similar Morphology to MEFs In Vitro

If iPSCs and MEFs demonstrate similar phenotypes for adipogenesis *in vitro*, we could begin by using a similar implantation approach. Morphology was evaluated and compared between PDiPSCs and MEFs grown in monolayer with expansion and adipogenic media. The PDiPSCs in adipogenic media demonstrated evidence of lipid containing cells by day 3, as seen by the positive ORO staining [Figure 5a], while the MEFs in adipogenic and expansion media conditions started to form lipid containing cells around day 5. As the number of PDiPSCs appeared to increase with time in adipogenic media, the same trend was not observed for the MEFs. The MEFs in adipogenic media generated similar numbers of lipid containing cells from days 7-14. Interestingly, the MEF expansion media condition did show some indication of lipid containing cells, although they appeared less phenotypically healthy and round compared to the adipogenic media condition. PDiPSCs and MEFs were stained with BODIPY to determine if the lipid containing cells were adipocytes [Figure 5b]. The BODIPY staining demonstrated adipogenesis in the MEF adipogenic media condition at around day 5, and the number of adipocytes appeared to increase throughout the 14-day time course. Similar trends were observed with the PDiPSC adipogenic media condition. Lipid content and differentiation efficiency were quantified from the BODIPY confocal images to understand if there were any differences between MEFs and PDiPSCs grown *in vitro* [Figure 5c]. With respect to the percent of lipid content at day 14, both the PDiPSC (38.7 ± 2.64%, *p<0.*001) and MEF (16.7 ± 6.62%, *p<0.*003) groups grown in adipogenic media had significantly increased percentages of lipid content compared to the PDiPSC expansion control group. The PDiPSC adipogenic media condition had a significantly increased differentiation efficiency (0.52 ± 0.15, *p<0.*001) compared to the control group, while the MEF adipogenic group did not (0.15 ± 0.01, *p<0.166*). MEFs and PDiPSCs were evaluated for adipogenic gene expression over time and were compared to PDiPSCs in expansion media as the control. mRNA levels for *Ppar*γ in both MEF groups grown in expansion and adipogenic media had significantly higher gene expression compared to the control (expansion media *p<0.001,* adipogenic media *p<0.005*) [Figure 5d]. However, these levels were reduced compared to the PDiPSC adipogenic media condition (*p<0.001*). *Apn* expression was analyzed, and neither MEF group was significantly different from the control, unlike the PDiPSC adipogenic media condition (*p<0.001*). Lastly, *Lep* expression was also different between the cell types. MEF groups demonstrated significantly higher mRNA levels for *Lep (*expansion media *p<0.005,* adipogenic media *p<0.016*), but these levels were still lower than the PDiPSC adipogenic group (*p<0.001*).

**Fig. 5.**
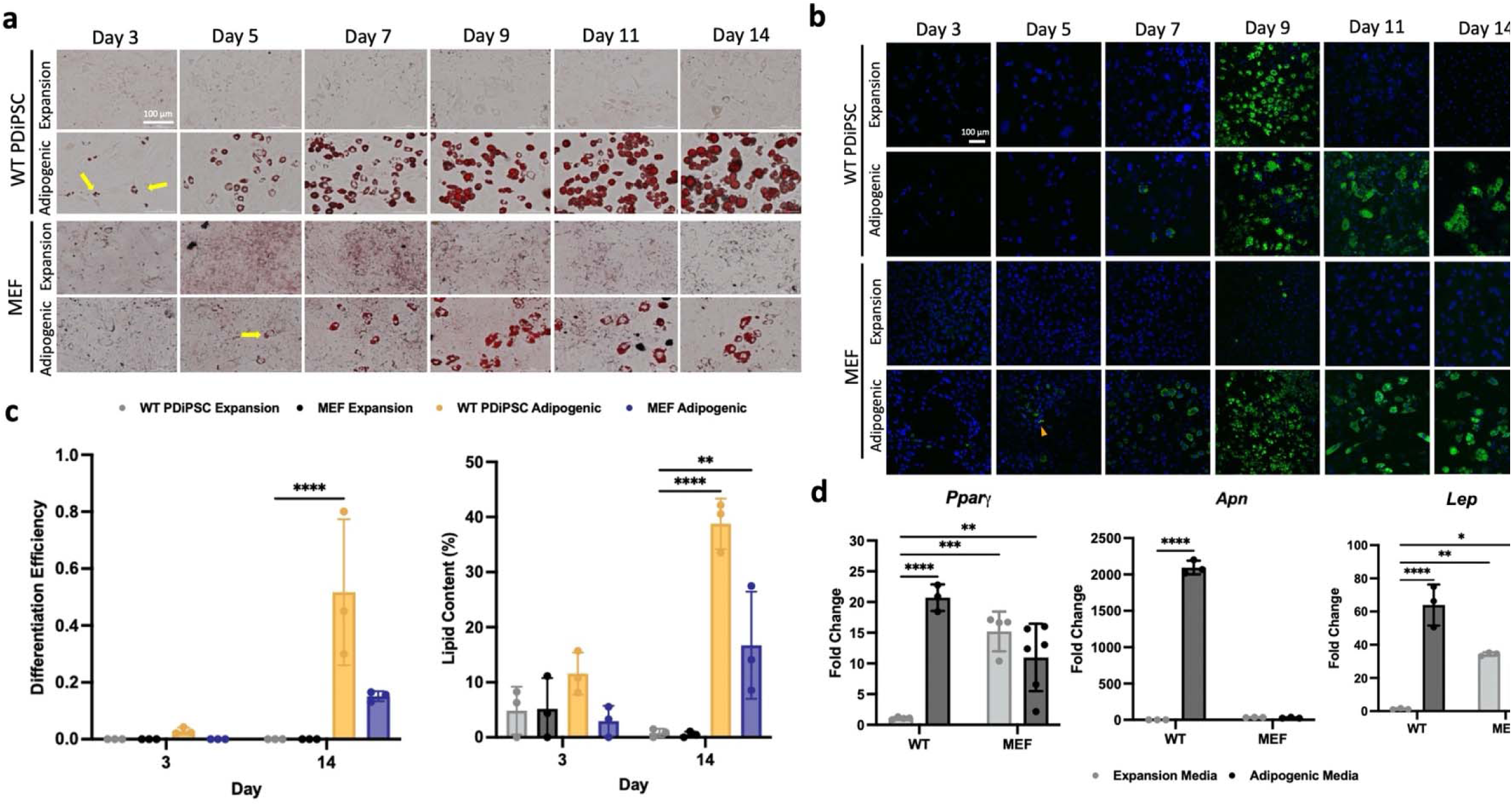
PDiPSCs and MEFs display similar adipogenic morphology in culture. WT PDiPSCs and MEFs were cultured in expansion or adipogenic media for 14 days, (a) groups were stained with Oil Red O stain, where the red stain indicates lipid content in cells. The yellow arrows point to lipid-containing cells. Scale bar is 100pm. (b) groups were stained with BODIPY/DAPI at various timepoints. BODIPY, green, stains for lipids and DAPI, blue, stains nuclei. Scale bar is 100pm. (c) Differentia efficiency and lipid content of all groups was measured at days 3 and 14 using the BODIPY images. Bars represent differentiation efficiency or lipid content (%) ± S (n=3). *Asteriks* represent significance (** *p<.005,* **** p<.00005) compared with the PDiPSCs expansion media control, (d) Cells from all groups above were collected at various timepoints for gene expression characterization, day 9 is shown here. *Ppgr?, Adip,* and *Lep* were target genes evaluated. Bars represent fold change ± SEM (n=3). Let represent significance *(A p<.05, PDiPSCs adipogenic media; B p<.05 MEFs expansion media; C p<05 MEFs adipogenic media).* All statistics were run using a 2-way ANOVA ANOVA with Sidak’s post-hoc test. SEM, standard error of the mean.

### Designer Adipocytes Engraft and are Functional In Vivo for 28 weeks

To image and measure cell signaling over time, we utilized Ccl2-Luciferase iPSCs (Ccl2 Luc), which were engineered to produce luciferase under the endogenous Ccl2-promoter in the presence of low-level inflammation (Brunger *et al*., 2017b; Hao and Baltimore, 2009). Mice were injected with a PBS control, 10e^6^, or 20e^6^ adipocyte media primed Ccl2-Luciferase PDiPSCs and then imaged using IVIS imaging [Figure 6a]. Mice given the higher cell dose, 20e^6^, had a consistently higher photon flux compared to the mice given 10e^6^ cells and were trending or statistically significant from control PBS injection at week 1 (*p<0.059*) and week 2 (*p<0.031*) across the 4 weeks after injection. A longer-term *in vivo* implantation study was then used to determine the length of time the implanted designer adipocytes were detectable *in vivo*. One week after injection, localized luminescent signal was present in the sternal region of WT and LD mice where the cells were injected [Figure 6c]. Two weeks after injection, the WT mice exhibited a marked decrease in luminescence in the sternal region, but the LD mouse maintained robust concentrated signaling. Then, six weeks after injection, there was a considerable reduction in localized luminescence in the WT mouse, and robust signaling was maintained in the LD mouse. However, after 10 weeks, localized luminescence was lost in both groups. These findings were corroborated by calculating the photon flux of both groups along the time course [Figure 6b]. After ten weeks, WT mice photon flux levels did not exceed that of background luminescence, while LD luminescence was reduced but detectable through 28 weeks.

**Fig. 6.**
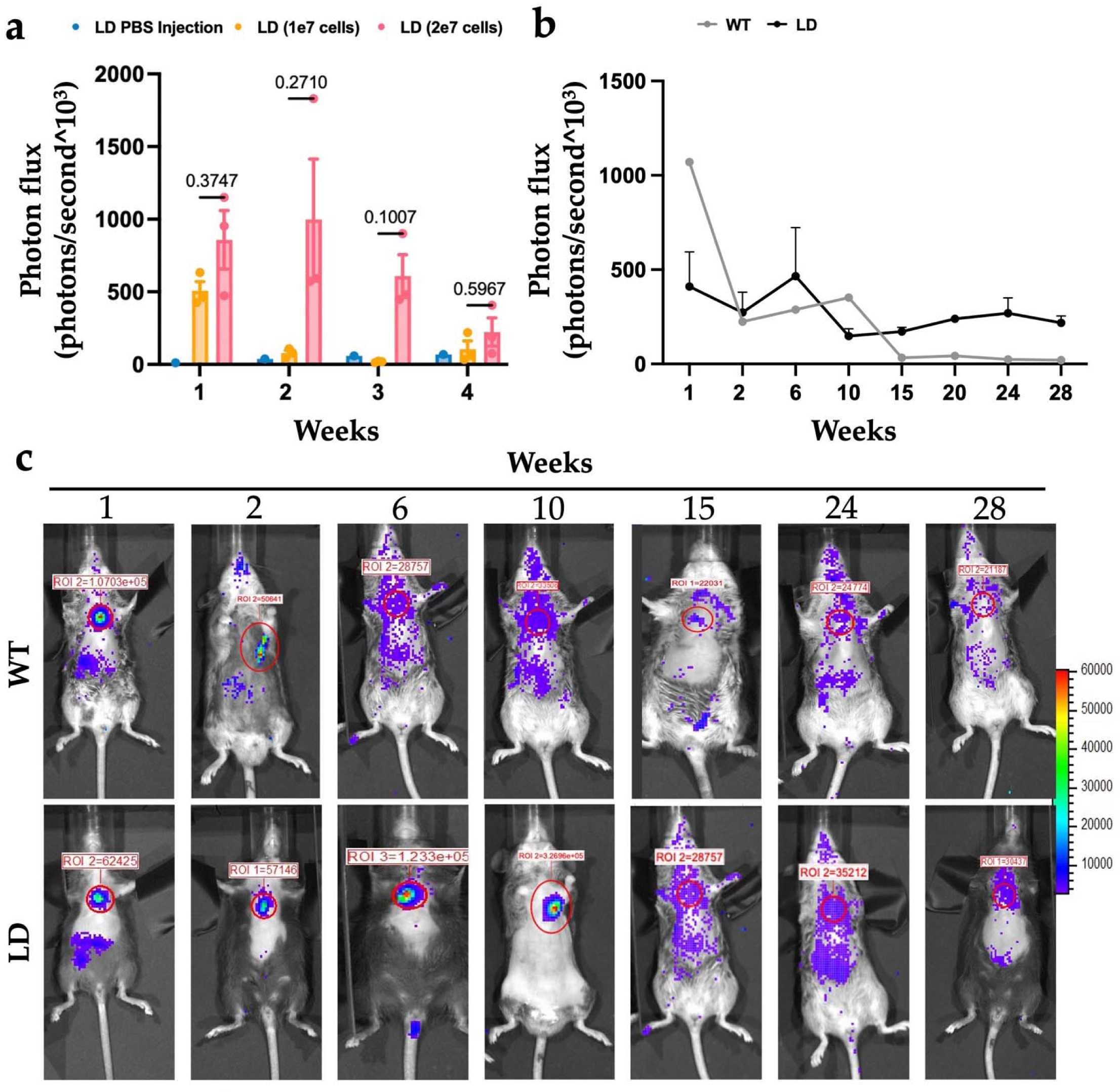
Self-regulating adipogenic iPSCs are viable and functional in vivo for up to 28 weeks. Different cell concentrations of Ccl2-Luciferase PDiPSCs primed in adipogenic media for 3 days were injected into LD mice and IVIS imaging was used to quantify cells present in the mouse, (a) Photon flux per animal was calculated every week for 4 weeks. Bars represent mean photon flux + SEM *(n=3-* 7). Exact *p values* are displayed using LD mice with PBS injection as the control, (b) WT and LD mice were injected with 20e^6^ cells and imaged for 28 weeks using IVIS imaging. Values represent mean photon flux ± SEM *(n=3-7).* All statistics were run using a 2-way ANOVA with Sidak’s post-hoc test. SEM, standard error of the mean, (c) Representative images of WT and LD mice at various timepoints from IVIS imaging are shown. The red circle highlights the area of injection.

## Discussion

In this study, designer adipocytes were created that can both individually dissect the role of adipokines in a variety of contexts and harness these cells to deliver anti-cytokine therapies. Importantly, this work demonstrated that adipocytes can be derived from murine iPSCs in a virus-free manner. CRISPR-Cas9 genome editing was then utilized to generate designer adipocytes without leptin, illustrating that leptin is required for iPSC-adipogenesis. Leptin addition to culture media rescued the loss of adipogenesis and allowed for the growth of leptin KO adipose (Scheller *et al*., 2010). We then created a self-regulating adipocyte drug delivery system and confirmed that these iPSCs (Brunger *et al*., 2017a; Brunger *et al*., 2017b) can differentiate into adipocytes and produce similar drug levels compared to other differentiation protocols. We also derived white and brown adipocytes using a straightforward approach for iPSC differentiation. Lastly, we demonstrated that the designer adipocytes can be implanted into fat-free mice for a functional *in vivo* readout of cytokine signaling from adipose tissue.

Developing a non-viral protocol for iPSC adipogenesis was an essential focus of this work to minimize the number of modifications or perturbations with which the cells would be challenged. Given the goal of using CRISPR-Cas9 gene editing, which involves guide RNAs that may be delivered using viral vectors to knock out adipokines of interest, reducing the number of transfections the cells would experience was critical. Our data suggest that the virus-free adipogenesis protocol recapitulated morphology, lipid content, and adipogenic gene expression of well-known differentiation methods, and thus, we determined that iPSCs do not require a viral overexpression of *Ppar*γ to achieve robust adipogenesis. This simplified protocol facilitates the use of iPSCs as a renewable source for adipocytes.

Designer adipocytes can be derived from iPSCs with a single gene knockout and used to disentangle specific adipokines’ roles in various disease states directly. We validated this tool by first knocking out leptin in iPSCs, which once differentiated into adipocytes, can be used to directly unravel the role of leptin signaling in disease progression *in vivo,* independent of body weight and metabolism. Leptin is an adipokine and pro-inflammatory mediator that is increased with obesity and posited as a mediator for several diseases like cardiovascular disease (Kang *et al*., 2020) rheumatoid arthritis (Mounessa *et al*., 2016) and is among the most widely implicated factors in OA pathogenesis (Griffin and Guilak, 2008; Griffin *et al*., 2009; Yan *et al*., 2018). While culturing the leptin knockout PDiPSCs using the validated adipogenesis protocol, we confirmed that leptin is an important regulator and inductor of adipogenesis *in vitro,* consistent with previous reports (Palhinha *et al*., 2019; Tencerova *et al*., 2019; Yue *et al*., 2016), and in this case, was required for PDiPSC adipogenesis. We observed that the leptin KO PDiPSCs grew in an uncontrolled manner and, at times, were overconfluent and lifted from the well when using typical cell seeding conditions. To rescue adipogenesis, we added murine leptin at physiological levels similar to those previously reported (Picó *et al*., 2022; Wagoner *et al*., 2006) to the cell culture media and were able to recover morphology, increase gene expression profiles, and importantly, we created a unique platform to individually test the mechanism of deleting various adipokines using genome engineering. Coupled with immuno-compromised or fat-free mice, these designer adipocytes enable researchers to directly test the mechanistic role of leptin-adipose signaling *in vivo* in a well-controlled manner.

To generate an adipocyte-based drug delivery strategy using iPSCs, we validated that Ccl2-IL1Ra PDiPSCs, previously created by Brunger et al. (Brunger *et al*., 2017a; Brunger *et al*., 2017b), were capable of adipogenesis. Previously, these cells were differentiated into a chondrocyte-like phenotype to generate living anti-cytokine implants (Choi *et al*., 2021; Collins *et al*., 2023). The Ccl2-IL1Ra-derived adipocytes had similar morphological characterization and differentiation efficiencies and, importantly, maintained the ability to produce IL-1Ra to similar levels as those differentiated with expansion media in the presence of low-level inflammation. Of interest, we previously found that using a cell’s intrinsic promotor to transcribe and ultimately deliver IL-1Ra resulted in similar levels to constitutive delivery yet yielded better disease mitigation. This finding not only illustrated intriguing therapeutic potential for cells, but also suggested that developing cell-based drug delivery strategies of known mediators and to accommodate new mediators as they are discovered (Choi *et al*., 2021). Overall, these findings outlined a new cell-based anti-cytokine therapeutic approach that can be used to combat the low-level inflammatory environment propagated by obesity, OA (Haseeb and Haqqi, 2013; Robinson *et al*., 2016), and several other conditions.

A simplified approach to generating brown and white iPSC-derived adipocytes was explored in this work. Not all adipose tissue functions similarly, and it is crucial to consider the various subtypes of adipose and their implications on metabolism, regeneration, and overall health, especially given rich interest of harnessing the therapeutic potential of brown adipose in obesity (Bryniarski and Meyer, 2019; Lidell *et al*., 2014; Payab *et al*., 2021; Trayhurn, 2018; White *et al*., 2019), musculoskeletal applications such as rheumatoid arthritis (Saito, 2013; Singh *et al*., 2021) and skeletal muscle regeneration (Bryniarski and Meyer, 2019), and non-musculoskeletal applications like type 2 diabetes mellitus (Ailhaud *et al*., 1992; Lee *et al*., 2010). These data demonstrated that different media formulations (Obatala Sciences) are sufficient to induce brown or white adipogenesis using PDiPSCs from the morphological characterization and differentiation efficiency analysis. While we expected to see significant differences in lipid droplet size between the white and brown PDiPSC-derived adipocytes, as these are defining characteristics that distinguish the two, there was not a substantial difference in lipid droplet size between the two in these studies. This lack of a difference in lipid droplet size could be due to the timepoint we analyzed and is the focus of ongoing work. Significant changes in adipogenic gene expression were only observed for adiponectin, while *Pparγ* and leptin were not significant by day 7 in culture. Importantly, *Ucp-1*, a marker for brown adipocytes, was increased compared to the control. Perhaps at an early timepoint of 7 days, PDiPSCs can likely be successfully differentiated into brown and white adipocytes, but the culture must be analyzed through day 14 to see apparent phenotypic differences in the outcomes we assessed. The three depots of adipose tissue – white, brown, and beige – are phenotypically and functionally distinct (Lidell *et al*., 2014; Liu *et al*., 2022; Soler-Vázquez *et al*., 2018; Trayhurn, 2018), and the ability to engineer these distinct adipose tissues and individually knockout genes of interest can help us to understand potential therapeutic or harmful effects of different adipose subtypes. Previous groups have successfully generated brown-like ADMSCs using genome engineering techniques and the beneficial effects of transplantation of these cells into obese mice, including improved glucose tolerance, insulin sensitivity, and increased energy expenditure (Wang *et al*., 2020). This efficient approach of generating diverse types of adipose tissue from iPSCs is viable, and it will allow future applications to integrate genome editing with these tissues, creating new possibilities for cell-based therapeutics, as previously explored with adipose-derived stem cells (Lopez-Yus *et al*., 2023). By simplifying the downstream differentiation of these three phenotypes of adipocytes, we can then use genome engineering to knock out genes of interest or rewire the different adipocyte depots for drug delivery applications.

To enable translational use of these designer adipocytes, we demonstrated that these cells can engraft and survive after implantation *in vivo* for up to 28 weeks. For these studies, we aimed to recapitulate the methods used to inject mouse embryonic fibroblasts subcutaneously into mice, as these proved successful in creating a fat depot 21 days after injection (Brenot *et al*., 2020; Collins *et al*., 2021; Ferguson *et al*., 2018). We utilized Ccl2-Luciferase PDiPSCs (Brunger *et al*., 2017a; Brunger *et al*., 2017b) primed with adipogenic media in culture for three days before injection into mice. We then tested the feasibility of using these implanted cells in a longer-term 28-week study. Detectable luciferase signal from the Ccl2-Luciferase cells was observed over the 28 weeks, concordant with our lab’s previous data (Collins *et al*., 2023); however, after six weeks, localized luminescence was lost at the sternal region where the cells were injected. These data highlight the feasibility of implanting these designer adipocytes *in vivo*, however, more work must be done to enhance this approach and cell survival in the mouse.

While this study represents several advancements in the potential use of iPSCs to engineer adipocytes, it is not without limitations. First, when characterizing and evaluating the feasibility of creating distinct depots of adipocytes, there was a limited supply of media, so the 14-day time course used in other sections was abbreviated. In the abbreviated culture in this study, brown and white adipocytes were successfully differentiated using the media; however, the lack of significant findings in the gene expression characterization are likely due to the shortened culture timeline. Additionally, initial work in this study to transplant the designer adipocytes into mice highlights the need for a refined approach in future studies to increase localized cell engraftment, which is essential in various transplantation applications. The methodology from MEF transplantation was used and compared using PDiPSCs. There are differences that may point to why the MEFs are able to engraft into the host and why the PDiPSCs may disperse. MEFs are a heterogeneous “pre-differentiated” population where PDiPSCs have undergone an MSC-like differentiation, and while they are heterogeneous, it is unknown which cell types are evident and how they facilitate the generation of the fat pad or immune privilege of the cells that allow for fat pad growth. Ongoing work will profile MEFs to determine if a population of cells primed for engrafting exists that is not present in the PDiPSCs. To facilitate engraftment, ongoing work is focused on using biomaterials or 3D constructs that can work synergistically with the designer adipocytes to help maintain them in the mouse (Yang *et al*., 2017), like Poly (ethylene glycol)-diacrylate (PEGDA) hydrogels or 3D woven scaffolds (Choi *et al*., 2021). Future steps for the designer gene knockout adipocytes will include creating more designer adipocyte knockout lines, including adipokines such as visfatin, resistin, and adipsin.

## Conclusion

The development of iPSC-derived adipocytes, or designer adipocytes, offers a unique opportunity to unravel the intricate roles of adipokines in health and disease progression. Through our research, we have demonstrated the potential of engineered adipose tissue as a novel drug delivery strategy and explored diverse designer adipocyte populations, including adipokine/gene knockout adipocytes, drug delivery adipocytes, and brown and white adipocytes. Despite the proof of concept demonstrated through the *in vivo* implantation studies in mice, there remains a need for a refined delivery approach, possibly through the integration of biomaterials or 3D scaffolds.

This work establishes a systematic framework for dissecting the functions of adipokines in disease progression, utilizing reprogrammed adipocytes as therapeutic depots and investigating the multifaceted roles of various adipose subtypes in disease and tissue therapeutics. Our controlled system, engineered from single cell isolates and implantable into fat-free mice for functional assessments, addresses many existing limitations in studying the downstream consequences of adipose signaling. The ultimate translational goal of this research is to employ the designer adipocytes developed in this study to meticulously dissect the involvement of adipokines in interorgan crosstalk, opening new avenues for understanding and targeting obesity-related diseases.

## Methods

### PDiPSCs Isolation and Culture

Murine induced pluripotent stem cells (iPSCs) were isolated and cultured from tail fibroblasts of adult C57BL/6 mice that were reprogrammed through forced overexpression of *Oct4, Sox2*, *Klf4*, and *c-Myc* as reported previously (Diekman *et al*., 2012). In the current study, four different lines of iPSCs were used: unedited/wildtype (WT), leptin (*Lep*^-/-^) knockout (described below), Ccl2-Luciferase (Ccl2-Luc), and Ccl2-IL1Ra. Ccl2-Luc and Ccl2-IL-1Ra iPSCs were previously created in the lab by incorporating luciferase or IL-1Ra under the endogenous *Ccl2* promoter (Brunger *et al*., 2017a; Brunger *et al*., 2017b; Pferdehirt *et al*., 2022). iPSCs were then differentiated into post-differentiated iPSCs (PDiPSCs) through a high-density micromass culture, as previously described (Diekman *et al*., 2012), which directs them towards a mesenchymal state. From this procedure, PDiPSCs were attained and either frozen at passage 1 or maintained through adipogenic or expansion media culture.

### Monolayer Adipogenesis Culture

To create a streamlined protocol for PDiPSC-adipogenic differentiation, two methods were compared: 1) overexpression of *Ppar*γ using lentivirus and 2) the use of commercially available adipogenic media for MSCs. The virus used in this experiment was pMSCV-Pparγ virus, which was previously constructed and described (Tontonoz *et al*., 1994). For these comparisons, PDiPSCs were first plated on gelatin coated 6-well plates (150239; Thermo Fisher) or 8-well cell culture slides (CCS-8; Mattek) for 1 day at a density of 4.16e^4^ cells/cm^2^ (4.16e^5^ cells/well) in expansion media consisting of high glucose Dulbecco’s modified Eagle’s medium (11-995-081; Gibco), fetal bovine serum (S11550; Atlanta Biologicals lot #A17004), 2-mercaptoethanol (21-985-023; Fisher Scientific), ITS+ premix (AR014; R&D Systems), penicillin-streptomycin (P4333-100ML; Sigma-Aldrich), L-ascorbic acid 2-phosphate (50μg/ml; A8960-5G; Sigma-Aldrich), L-proline (40μg/ml; P5607-25G; Sigma-Aldrich), and transforming growth factor–β3 (10 ng/ml; 243-B3-200; R&D Systems). On day 1 of culture, cells were transduced with equal titers of virus as previously described (Tontonoz *et al*., 1994), where titers were described as doses. Cells received either a low dose (3.1µl/cm^2^), a high dose (5.2µl/cm^2^), or no virus in 2.5mL polybrene-containing expansion media without ascorbate and proline. On day 2, all wells were washed 2 times with growth media. On day 3, wells were puromycin-selected with 2.5µg/mL of puromycin. On day 4, wells received either expansion media commercially available adipogenic media (Mesencult Adipogenic Differentiation Kit; 05505; STEMCELL). Media was changed and cells were collected for analysis every other day up to day 14. The standard culture conditions referred to in the subsequent methods are PDiPSCs cultured in expansion or adipogenic media, with no virus.

### Generation of Leptin Knockout iPSCs

Leptin knockout (KO) iPSCs were created utilizing resources from the Genome Engineering & iPSC Center at Washington University in St. Louis. Using CRISPR-Cas9, two guide RNAs (YH665.m.Lep.sp7 and YH666.m.Lep.sp9) were selected based on an off-target analysis and distance to the target site as seen below. Murine leptin has 2 exons and thus a deletion approach was used to complete the knockout.

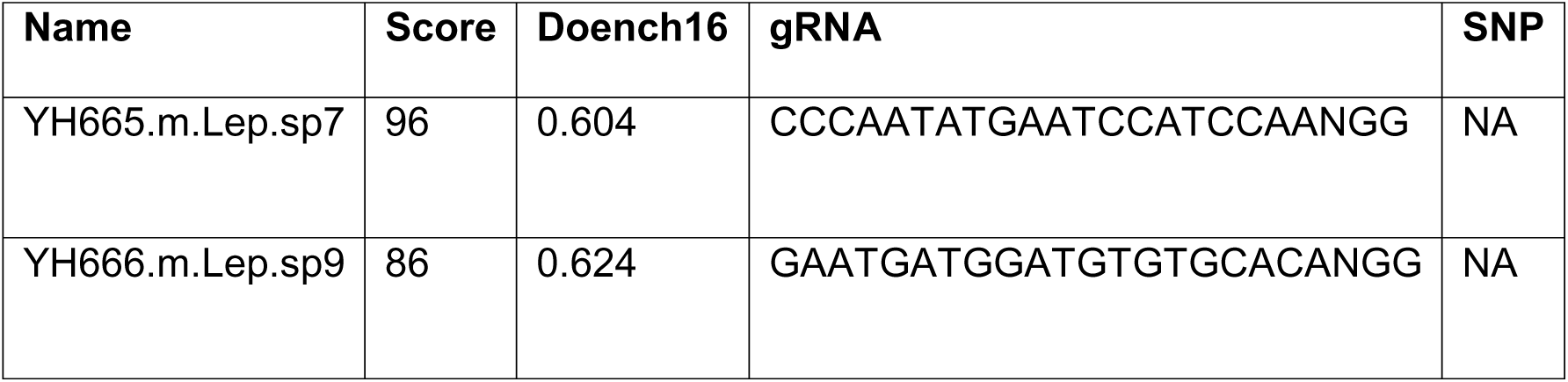

Score was computed as 100% minus a weighted sum of off-target hit-scores in the target genome (full ranked list not shown). SNP NA indicates no common SNP found in the gRNA. Leptin KO iPSCs were then differentiated into PDiPSCs using the micromass procedure as described above.

### Leptin Supplement for Adipogenesis Culture

In culture, the leptin knockout PDiPSCs received doses of leptin to test whether loss of adipogenesis can be rescued under standard culture conditions. Leptin knockout PDiPSCs in culture were supplemented with 1,000ng/mL of murine leptin (498-OB-05M; R&D Systems), a concentration that was determined after performing a leptin dose analysis [Figure 2i]. The literature supports doses anywhere from 250-1,000ng/mL (Picó *et al*., 2022; Wagoner *et al*., 2006). Cells received this dosage with each media change from the time they were plated until the end of the adipogenesis time course.

### Brown and White Adipocyte Culture

Various adipogenic media were assessed for their ability to efficiently generate distinct adipocyte sub-populations from iPSCs in monolayer. WT PDiPSCs were plated at a density of 4.16e^4^ cells/cm^2^. Cells were incubated for 24 hours in 5% CO2 at 37 °C, then fed with iPSC expansion media (described above) thereafter until 80-90% confluency was reached (1-2 days). Upon exhibiting the desired confluency, PDiPSCs were cultured in the presence of the different fat-specific media types. Cells were maintained in either a PDiPSC expansion media control or one of the following commercially available adipogenesis media: STEMCELL Mesencult MSC Basal Medium with adipogenic supplement (Mesencult Adipogenic Differentiation Kit; 05505; STEMCELL) (adipogenic media), AdipoQual BAT Medium (OS-013; Obatala Sciences) (brown media), or Obatala AdipoQual WAT Medium (OS-014; Obatala Sciences) (white media). Cells were cultured in their respective media for up to 7 days of adipogenesis, with media changes taking place every other day. Sample collections occurred at days 0, 1, 3, 5, and 7 for staining and imaging.

### Mouse Embryonic Fibroblast Culture

Mouse embryonic fibroblasts (MEFs) were used as a control cell line in this study as previous work demonstrated great success with implanting these cells into a mouse model (Collins *et al*., 2021; Ferguson *et al*., 2018). MEFs were prepared as previously described (Ferguson *et al*., 2018) and cultured at a density of 4.16e^4^ cells/cm^2^ in expansion or adipogenic media for up to 14 days.

### Gene Expression

Samples for gene expression analysis were harvested and total RNA was isolated from the cell according to recommendations of the manufacturer (48300; Norgen Biotek). Reverse transcription was performed using the superscript VILO complementary DNA synthesis kit (11755500; Life Technologies) following the manufacturer’s instructions. Quantitative polymerase chain reaction (qPCR) was performed using FASTSybr (4385617; Applied Biosystems) following manufacturer’s instructions. Cycling parameters were initial denaturation at 95°C for 10 minutes followed by 40 cycles of 95°C for 15 seconds and 60°C for annealing and extension for 60 seconds. Gene fold changes were determined relative to a control group using 18S ribosomal RNA as a reference gene. Data are reported as fold changes and were calculated using the 2^–ΔΔCt^ method. Primer pairs were synthesized by Integrated DNA Technologies, Inc.

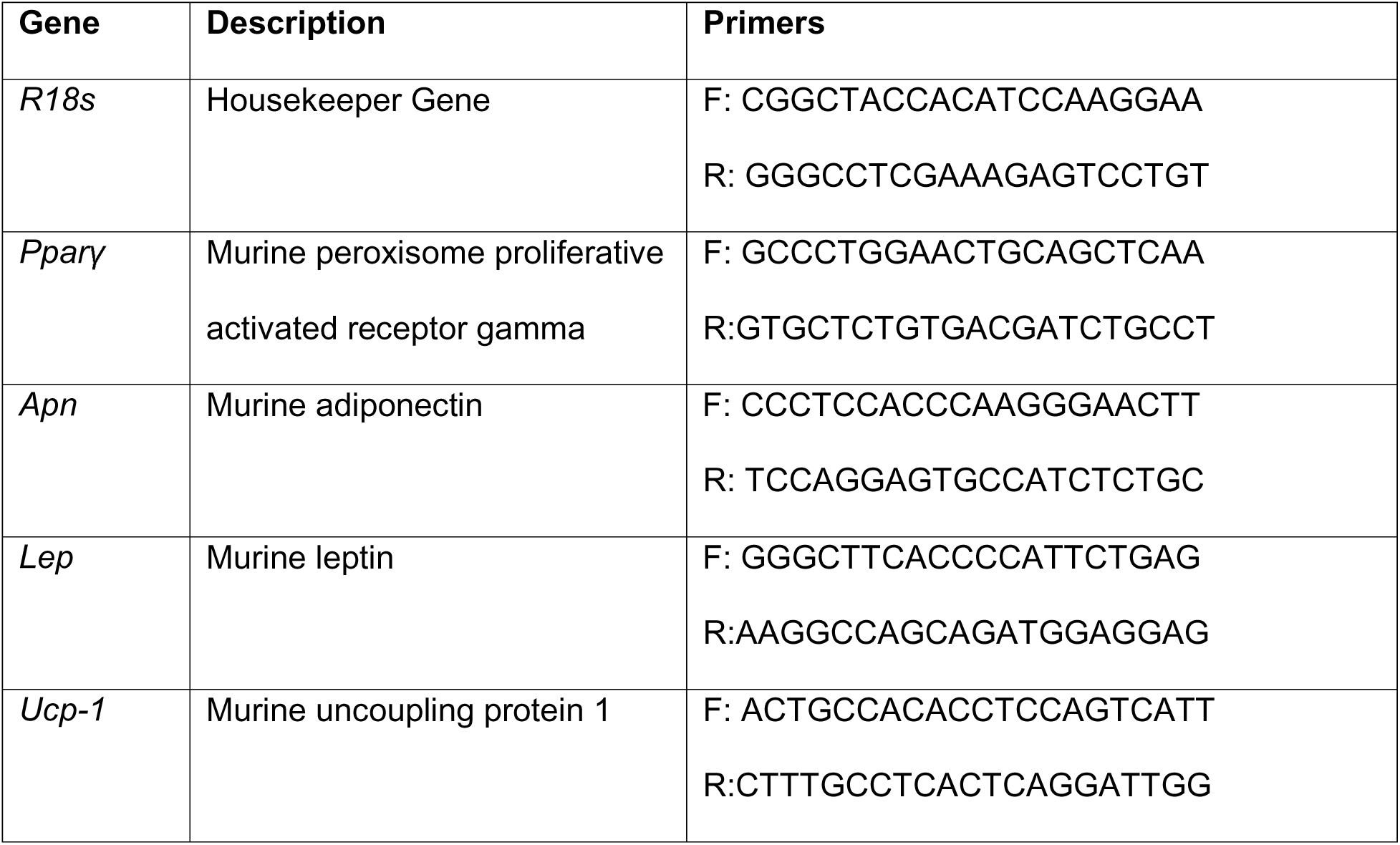

### Inflammatory Challenge

To measure the kinetics of IL-1Ra production in the Ccl2-IL1Ra PDiPSCs, WT and Ccl2-IL-1Ra PDiPSCs were treated with expansion and adipogenesis media until day 14. On day 14, the cells were challenged with 1 ng/mL murine IL-1α (400ML005CF; Fisher Scientific). 500µL of culture media was collected and snap frozen at 0, 4, 12, and 72 hours post inflammatory challenge. Control groups were not challenged with IL-1α.

### Enzyme-linked immunosorbent assays

Enzyme-linked immunosorbent assay (ELISA) was used to measure the concentration of IL-1ra *in vitro* using a mouse IL-1Ra DuoSet ELISA kit (DY480; R&D Systems) with a 1:10 dilution. Each sample was measured in technical duplicates and absorbance was measured at 450 and 540nm (n=4-6 per group). The lower limit of detection for the ELISA was 156 pg/mL.

## Histological Characterization

### Oil Red O Staining

Oil Red O is a stain for neutral lipids and is used to assess adipocyte differentiation. All lines of PDiPSCs were characterized qualitatively by Oil Red O staining to assess morphology and lipid content. A stock solution was created by adding 0.5g Oil Red O (O0625-25G; Sigma-Aldrich) to 100mL isopropanol. A working solution was curated from the stock solution using 3 parts stock solution and 2 parts deionized (DI) water. Cells were fixed with 4% paraformaldehyde for 15 minutes. Paraformaldehyde was then aspirated and then 1mL of working solution was added to the each well of a 6-well plate to cover the cells for 30 minutes. Working solution was then aspirated and the wells were washed 1x with DI water. DI water was added back into the wells and cells were imaged using a Cytation 5 (Biotek; Agilent, Santa Clara, California).

### BODIPY Stain

PDiPSCs cultured in 8-well cell culture slides (CCS-8; Mattek) were stained with BODIPY 493/503 (4,4-Difluoro-1,3,5,7,8-Pentamethyl-4-Bora-3a,4a-Diaza-*s*-Indacene) (D3922; Invitrogen), a green, fluorescent dye that stains lipids, and a DAPI (4’,6-Diamidino-2-Phenylindole, Dilactate) (422801; BioLegend) counterstain. Briefly, cells were fixed with 4% paraformaldehyde for 5 minutes and then washed 2x with PBS. A 1:200 BODIPY solution was added to each well for 20 minutes. The wells were then washed 3x with PBS. A 1:1,000 DAPI solution was added to each well for 7.5 minutes. The wells were then washed 3x with PBS and then the slide was cover slipped using ProLong Glass Antifade Mountant (P36982; Invitrogen). Slides were then imaged using confocal microscopy (LSM 880, Zeiss, Thornwood, NY, USA).

## Image Analysis

### Quantifying Differentiation Efficiency

A custom ImageJ workflow was developed to quantify adipocyte differentiation efficiency from PDiPSCs and MEFs during adipogenesis. First, the total number of adipocytes were manually counted from 20x BODIPY/DAPI images. Cells were deemed adipocytes if they showed positive BODIPY staining surrounding DAPI-labeled nuclei. For nuclei surrounded by low signal, the corresponding cells were considered adipocytes if adjacent regions contained distinct, circular lipid droplets, and not hazy, irregularly shaped, or low signal. To obtain an overall cell count, the DAPI image channel was first pre-processed for nuclei segmentation. Pre-processing steps included subtracting a value of 50 pixels across the entire image background. A low-pass filter was then applied to blur the image (Gaussian blur) with a sigma value of 4. An ImageJ plugin (StarDist (Schmidt *et al*., 2018)) was then utilized to measure the total number of nuclei in the resulting image. Finally, the differentiation efficiency was calculated by dividing the number of differentiated adipocytes by the total number of cells in an image (n=3 images/condition).

### Quantifying Lipid Droplet Size

A separate ImageJ workflow was developed for the quantification of lipid droplet content and size among PDiPSCs and MEF cells under various conditions undergoing adipogenesis. Briefly, 20x confocal images containing only the BOPIDY channel were first pre-processed for lipid droplet segmentation by converting to image type 16-bit and subtracting up to a 50-pixel radius across the entire image background to correct for hazy signaling due to cellular autofluorescence. The background-subtracted image was duplicated, after which a low-pass filter was applied to blur the duplicate image (Gaussian blur, σ = 4) and smooth intensity variations across the image. The blurred image was subtracted from the non-blurred image. The result of this operation was converted to a 32-bit image type for manual threshold application. A manual threshold value of 3 was applied to yield a binary image of segmented lipid droplets. To quantify lipid droplet size for the depot-specific adipogenesis experiments, a particle analysis was performed across the full image, yielding individual particle areas. The average of these area measurements was found for the image. This process was repeated for at least three images of the same condition.

### Quantifying Lipid Droplet Content

To quantify lipid droplet content, 5 cells were randomly selected from the binary image for further analysis. Target cells were selected by first utilizing the corresponding DAPI-only image as a reference to choose a single nucleus in one of five image quadrants (top left/right, bottom left/right, center). Then, the corresponding cell of interest was located in the binary image by referencing common target cell features between both the DAPI-only image and the multi-channel image. Upon locating target cell droplets in the binary image, the total areas of these particles were measured by tracing around the lipid-containing region of interest and enacting the particle analysis function. The perimeter of the target cell was then manually traced in the multi-channel image and total cell area was measured. For each of the 5 selected cells, the total lipid content was calculated by dividing the sum of the particle areas by the total area of the cell. The average lipid content was then determined across the 5 cells of the same image. This process was repeated for at least three images of the same condition.

### In Vivo Cell Delivery

To perform observations of cell delivery over time, we used fat free lipodystrophic (LD) mice (adiponectin-DTA) (Wu *et al*., 2018) characterized by a low-level systemic inflammatory environment that can drive the Ccl2-Luciferase circuit for *in vivo* imaging (Brunger *et al*., 2017b; Collins *et al*., 2021). First, to understand the concentration of cells to inject into the mice, Ccl2-Luc PDiPSCs were cultured until day 3 in adipogenic media then removed from tissue culture plastic using trypsin for 10 minutes to help remove excess matrix. Then, cells were centrifuged for 3 minutes at 1500rpm (500-600g) and counted. Cells were resuspended to yield final concentrations of either 10 or 20 million cells in 250μL sterile PBS for injection. Control LD mice received injections of 250μL PBS (n=3). LD mice were anesthetized using 2-3% isoflurane. Using a 27-gauge insulin syringe, cells were injected just superficial to the sternum slowly until the contents of the injection were delivered to the mice. Once the ideal cell number was determined (20e^6^ cells per injection), we performed a 28-week injection experiment using LD and WT mice. All animal protocols were approved by the Washington University School of Medicine IACUC under protocol 19-0774.

### In Vivo Imaging

Mice were prepared for imaging using Veet hair removal and standard clippers. For the cell number experiments, LD mice were imaged weekly for 4 weeks. Mice were imaged under 2-3% isoflurane anesthetic. D-luciferin substrate (150 mg/kg in phosphate-buffered saline (PBS); Gold Biotechnology, St. Louis, MO, USA) was delivered by the Molecular Imaging Core via intraperitoneal injection (IP). After waiting 10 minutes, mice were positioned on their dorsal aspect and images were taken from the ventral view with an exposure time of 5 minutes using an IVIS Lumina (PerkinElmer, Waltham, MA, USA; Living Image 4.2; 1-min exposure; bin, 8; field of view, 12.5 cm; f/stop, 1; open filter). Images were analyzed by the Molecular Imaging Core and a pre-defined region of interest (ROI) 1cm^2^ area superficial to the mouse sternum, operationalized as photon flux. Once the cell number experiment was optimized, a new subset of mice was imaged at 1, 2, 6, 9, 12, and 28 weeks timepoints.

### Statistics

Statistical analysis and sample numbers for each experiment are detailed in the respective figure legends. All statistics were performed in Graphpad Prism 8 (Graphpad Software, San Diego, Ca). Data are presented as means ± standard error of the mean.

## Acknowledgments

We thank Kristin Lenz for assistance with *in vivo* components of this study, Sara Oswald for laboratory support, the Genome Engineering & Stem Cell Center at WashU for assistance with genome editing applications, the Molecular Imaging Center at WashU for IVIS imaging, and Obatala Sciences for supplying media for the study. We thank Charles Harris for providing the virus, originally generated by the laboratory of Dr. Bruce Spiegelman. This work was funded by the Shriners Hospital for Children, the Washington University in St. Louis Institute of Clinical and Translational Sciences grant JIT693, and in part by NIH grants AR078949, AR080902, AR078949, AG15768, AG46927, AR072999, DK108742, AR073752, AR074992.

